# MegaKG: Toward an explainable knowledge graph for early drug development

**DOI:** 10.1101/2024.03.27.586981

**Authors:** Jianqiang Dong, Junwu Liu, Yifan Wei, Peilin Huang, Qiong Wu

**Affiliations:** MegaRobo Technologies Co., Ltd, Beijing, China

## Abstract

In biomedical research, the utilization of Knowledge Graph (KG) has proven valuable in gaining deep understanding of various processes. In this study, we constructed a comprehensive biomedical KG, named as MegaKG, by integrating a total of 23 primary data sources, which finally consisted of 188, 844 nodes/entities and 9, 165, 855 edges/relations after stringent data processing. Such a massive KG can not only provide a holistic view of the entities of interest, but also generate insightful hypotheses on unknown relations by applying AI computations. We focused on the interplay of the key elements in drug development, such as genes, diseases and drugs, and aimed to facilitate practical applications that could benefit early drug development in industries. More importantly, we placed much emphasis on the exploitability of the predictions generated by MegaKG. This may greatly help researchers to assess the feasibility or design appropriate downstream validation experiments, making AI techniques more than just black-box models. In this regard, NBFNet was adopted, which combines the advantages of both traditional path-based methods and more recently developed GNN-based ones. Performance evaluation experiments indicated superior results by MegaKG. We also conducted real case studies to validate its practical utility in various scenarios, including target prediction, indication extension and drug repurposing. All these experiments highlighted the potential of MegaKG as a valuable tool in driving innovation and accelerating drug development in pharmaceutical industry.

## INTRODUCTION

In the pharmaceutical industry, the driving force behind original innovation stems from a deep understanding of the molecular mechanisms that underlie disease biology. To date, a vast amount of discoveries have been made and are ever emerging in various aspects relevant in this field, encompassing molecular functions, cellular pathways, therapeutics, clinical phenotypes, and more. The interconnections among these entities are intricate and complex. Nonetheless, comprehending the relationships among these entities is pivotal in elucidating the mechanism of action of various processes. This, in particular, provides valuable insights into the causes and development of diseases and thus their possible therapies.

Decades of great effort has been made to systematically collect all these findings in certain forms of databases. Typically, these databases not only record information about a specific category of entities, but also capture the relations among multiple categories of entities. For example, DrugBank [1] saves drug-related relations, including drug-disease associations, drug-drug interactions, drug-gene targeting, and others. Starting from these primary high-quality biomedicine-related databases, more insights could be gained by integrating them into a knowledge graph (KG), i.e. a comprehensive heterogeneous multi-relational network of different categories of entities. Such a knowledge graph, like any specific graph in graph theory, is essentially a data structure composed of nodes and edges, with nodes representing the entities in biomedicine and edges the relations between the two connecting nodes (or entities). In this way, a holistic view can be provided, centered around the entities of interest, simply by extracting the interconnected collection of relevant entities and their relations. Furthermore, leveraging certain computational approaches on graphs enables the discovery of hidden relations which may not be immediately illuminated based on known facts alone. This holds great value in, for example, identifying potential therapeutic targets, exploring new avenues for drug discovery and treatment development, or other biomedical applications.

However, construction of such a biomedical KG is not quite straightforward. The knowledge is fragmented and scattered across various sorts of resources. A lack of consistency often exists in the terminology and naming conventions used for, for example, diseases or drugs. Some are organized using nested ontologies or represented at different scales, such as the pathway information. Even worse, many facts were accumulated in a non-standardized form, which still requires manual intervention from experts. Nonetheless, several biomedicine knowledge graphs have been developed, for different purposes. Reviews of these knowledge graphs can found in [2, 3]. For example, Hetionet was one of the first attempts toward a comprehensive KG suitable for various tasks [4]; Genetic and Rare Diseases Information Center (GARD) [5] focused exclusively on rare diseases, and advanced the understanding of under-diagnosed diseases; DRKG was developed upon Hetionet specifically for drug repurposing for Covid19 [6]; PharmKG was a compact one, focusing on entities of drug, gene and disease, and aimed for drug discovery [7]. The recently developed PrimeKG was specialized for precision medicine, and provided the most abundant nodes and relationships with dedicated annotations so far (129,375 nodes and 8,100,498 edges; see Table 1 for a comparison) [8]. However, there was a lack of inference on unknown relations, which greatly limited its application potential.

**Table 1:**
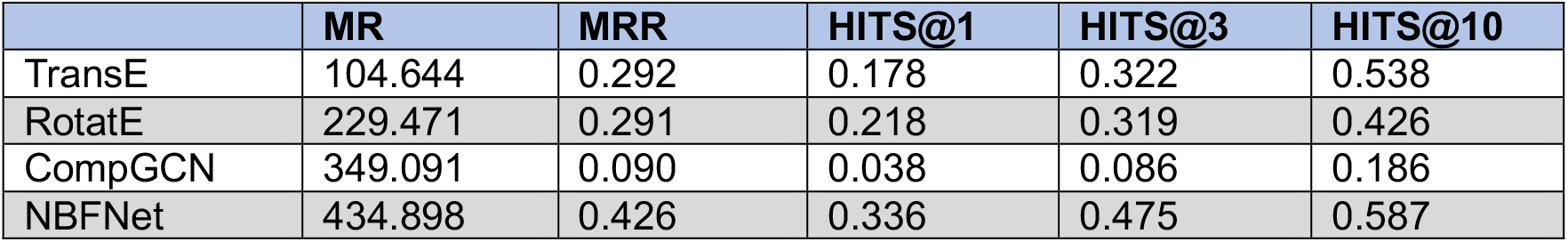
Performance comparison among representative models of different approaches.

The task of learning biological associations in this context can be modelled as link prediction in the form of (head entity, relation, entity) in knowledge graphs [9]. To maximize the predictive potential of a KG, it is essential to enhance the credibility of the predicted links. However, due to a lack of true negative samples, no proper benchmark dataset with ground truth information could be provided. Although metrics for evaluating the prediction accuracy of such positive facts enriched KG could be adopted [10], the persuasiveness is still limited. Especially in the pharmaceutical industry, the process of validating a hypothesis is usually lengthy and the cost of error may be rather high. Meantime, scientists also require valuable clues to design rigorous downstream experiments that can confirm or refute the hypothesis under investigation. In this context, the concept of explainability becomes crucial, which is also a prominent requirement for artificial intelligence to be applied effectively in various industries. In this regard, however, most of the KGs with inference capability do not offer explainability. One distinctive exception is PharmKG, which applied a graph attention network to assign a weight to each neighboring node of the given head entity. Although using such attention-based method could provide certain explainability, the way the attention PharmKG computed was static, namely, given a head node, the weights of its neighboring nodes were fixed, no matter what the tail node was. Alternatively speaking, the explainable basis stayed the same no matter what was asked to predict for a given head entity. Altogether, it is not difficult to see that improving the explainability of a KG is a imperative direction. This task becomes even more challenging when the size of the KG becomes rather large, which is often the case for a biomedical knowledge graph.

The dominant paradigm for such link prediction involves learning and operating on latent representations (i.e. embeddings) of entities and relations. Over the last decade, triplet-based models have gained much attention, which dissects a graph into a collection of triplets for each edge and its two associated nodes, and directly learn and reason on the embeddings of the entities and relations. Representative methods include TransE [11], ConvE [12], ComplEx [13], RotatE [14], etc. However, the triplets can only reserve the edge structure, which losts local connections. Meantime, triplet-based models are uninterpretable, and only transductive. Traditional path-based methods are inherently interpretable, since they can preserve sequential connections between nodes. Different streams of this type of methods have been developed including DeepPath [15], MINERVA [16], DRUM [17], PathCon [18], etc. However, these methods usually need to traverse an exponential number of paths and restrict to very short paths. Subgraphs can naturally capture more complex local information than paths. Graph neural network (GNN) based methods has been developed to capture the subgraph structures in KG. Representative methods including R-GCN [19], CompGCN [20] and KE-GCN [21] jointly embed both nodes and relations in the graph and update the representations of entities by aggregating all the neighbors in each layer. However, these GNN-based methods donot differentiate different neighbors and cannot provide interpretable predictions at all. Later attention based such methods [22], similarly with the HRGAT developed for PharmKG, can only provide aforementioned static interpretation.

Another stream of GNN-based frameworks, such as GraIL [23], was proposed to extract the enclosing subgraph between the head entity and tail entity. The key idea behind GraIL is to turn the prediction task into a logical induction problem. The learning process is entity-independent and thereby gains the inductive ability to generalize to unseen entities. Following GraIL, NBFNet [24] and RED-GNN [25] were proposed concurrently, with similar ideas to combine the advantages of both traditional path-based methods and more recently developed GNN-based ones. In this way, the link prediction task can be made interpretable, with inductive ability and high scalability.

In this study, we constructed a large biomedical knowledge graph, MegaKG, by integrating 23 high-quality resources, with ∼190, 000 nodes and ∼9 million relations (as shown in Figure 1). Orientated for early drug development and rooted for real needs from the pharmaceutical industry, MegaKG focused on the relevant entities of genes, diseases and drugs. NBFNet [24] was adopted to equip the KG with predictive, and more importantly, explainable capability. Centred around MegaKG, a series of direct applications can be achieved, including target discovery, drug repurposing, indication extension and drug interactions. Under a series of knowledge graph-specific assessments, NBFNet demonstrated superior performance compared to existing representative methods. We also provided several real cases that exhibit sound predictions with explainable path to derive the conclusions. We believe that MegaKG may serve as a powerful platform and engine for initiating projects in early drug development. We also anticipate the implementation of valuable insights derived from MegaKG within the industry.

**Figure 1.**
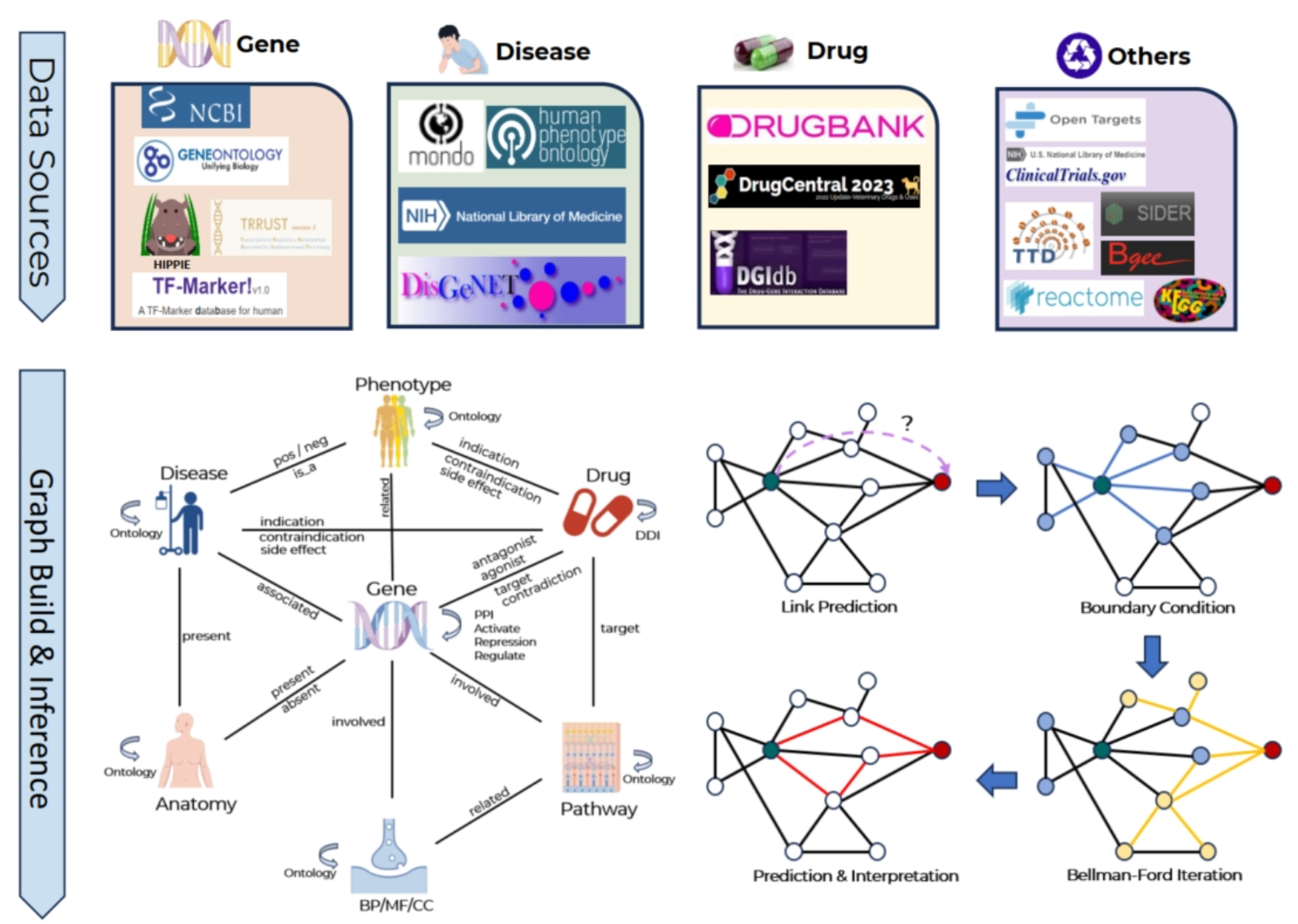
The overview of MegaKG

## MATERIAL AND METHODS

MegaKG was constructed using publicly available databases, with a focus on the fundamental elements that form the basis of early drug development, including genes, diseases and drugs. Stringent filtering criteria were applied to ensure the data quality underpinning the knowledge graph. Afterwards, relation prediction with interpretability was achieved by employing the NBFNet framework, which learns and reasons based on the subgraph between a head entity and a tail entity. The overview of MegaKG is shown in Figure 1.

In this section, we illustrate the details of the construction of MegaKG and the predictive model applied.

### Data sources to construct MegaKG

MegaKG was specifically designed to focus on the fundamental elements that underpin early drug development, including genes, diseases and drugs. Brief introduction to the relevant data sources are provided below. We tried to associate each database with gene, disease or drug-related categories, but please note that some databases may be relevant to multiple such categories.

#### Gene related databases

The Gene Ontology (GO) [26] defines the gene function hierarchies, where GO is organized into three main categories: Molecular Function (MF), Biological Process (BP) and Cellular Component (CC). NCBI GENE provides basic information of human genes and the associated GO annotations. KEGG [27] and Reactome [28] describes the pathway information highly relevant with gene functions. HIPPIE [??] focuses on protein-protein interactions and were incorporated by mapping the proteins to the corresponding gene entities. Databases specializing in certain elements playing important roles in regulating gene related processes, such as the transcriptional factors (TF-Marker [29] and TRRUST [30]), were also recruited.

#### Disease related databases

UMLS [31] and MONDO [32] describe the vocabulary and ontology of human diseases; HPO [33] is a database of phenotypic abnormalities pertaining to diseases. DisGeNET [34] (version 7.0) collects the genetic basis of human diseases. DAR[35] provides disease-pathway and disease-gene relations.

#### Drug related databases

DrugBank [1] and DrugCentral [36] are comprehensive platforms for drugs and drug-related entities. BioSNAP [37] contains drug-disease information. DGIdb [38] focuses on drug-gene interactions. SIDER [39] records adverse drug reactions. OpenTarget [40] and TTD [41] provide relationships between drugs, targets and diseases and serve as knowledge graphs of a small scale. ClinicalTrials [??] contains information of the systematic investigation of drug effects on human subjects.

#### Miscellaneous databases

For future development, we also include anatomy and gene expression information. UBERON[42] is a database of ontology of anatomical structures. Bgee[43] provides gene expression patterns across different tissues and organisms. Hetionet[4] was also incorporated to extract the disease-anatomy relationships.

### Data processing for integration

#### Entity normalization

A variety of organizations or projects have been dedicated to defining or annotating the genes, diseases or drugs. The first crucial step usually is to make the terminologies consistent, which can be a rather laborious process. What we did is as follow. Genes were aligned Ensembl ID/Gene Symbol/UniProt ID to Entrez Gene IDs based on NCBI [44]. For disease entities, we directly extracted reference from MONDO ontology or indirectly mapped UMLS CUI terms to MONDO followed PrimeKG[8]. Drug IDs were initially mapped to IDs provided by DrugBank. If no match was found, we attempted to map the ID to TTD. However, if neither mapping works, the drug term was excluded.

#### Entry filtering

For the disease-gene relations reported by DisGeNET [34], we only kept EI score equals 1, which indicates that all the publications support the gene-disease associations or the variant-disease associations; For the Bgee, similarly with PrimeKG [8], we set the selection criteria to be “expression_rank < 25000” and “call_quality == good quality”.

#### Overlapping relations across multiple sources

For a drug-gene relation, if the labels from different sources were the same, the unified label would follow as “agonist” or “antagonist” accordingly. Otherwise, if both of these two labels were found from different sources, we would mark the relation as “contradiction”. In other cases, if no indication on either label was provided, we just labelled the relation as “target”. For a drug-disease relation, if multiple records with different stage information were found, we just kept the most advanced one.

### Model construction

A knowledge graph is represented by G = (V, E, R), where V and E represent the set of entities (nodes) and relations (edges) respectively, and R denotes the set of relation types. For a given node *u*, we use N (*u*) to indicate the set of nodes connected to *u*, and E(*u*) the set of edges terminating at node *u*. Please note that for any node in V, there is always an edge connecting to itself (self-loop).

Link prediction on a knowledge graph focuses on predicting whether a query relation *q* exists between a head entity *u* and a tail entity *v*. In triplet-based methods, such as TransE [11] or RotatE [14], all the entities are independently trained by decomposing the graph in to a collection of (head, relation, tail) triplets, which may lose some reciprocal constraint embedded in the graph. To better capture the global relationships, a paired representation *h*_*q*_ (*u, v*), proposed by GraIL [23], that encapsulates the local subgraph structure between *u* and *v* with respect to the query relation *q*, is more appropriate.

Such a paired representation *h*_*q*_ (*u, v*) can be obtained by two steps, namely: 1. extracting a subgraph starting and ending at nodes *u* and *v*, respectively; 2. learning a representation on the subgraph.

To compute the subgraph corresponding to *h*_*q*_ (*u, v*), GraIL [23] identifies all nodes occurring on a path of length at most k between nodes u and v, and extracts the subgraph exclusively encompassing all these paths. For example, given a KG shown in Figure 2(a), with node ➀ as *u* and node ➄ as *v*, the subgraph to *h*_*q*_ (*u, v*) is displayed in Figure 2(b). NBFNet [24] and RED-GNN [25] similarly extract this subgraph, but with variations in their methods for computing paired representations.

**Figure 2.**
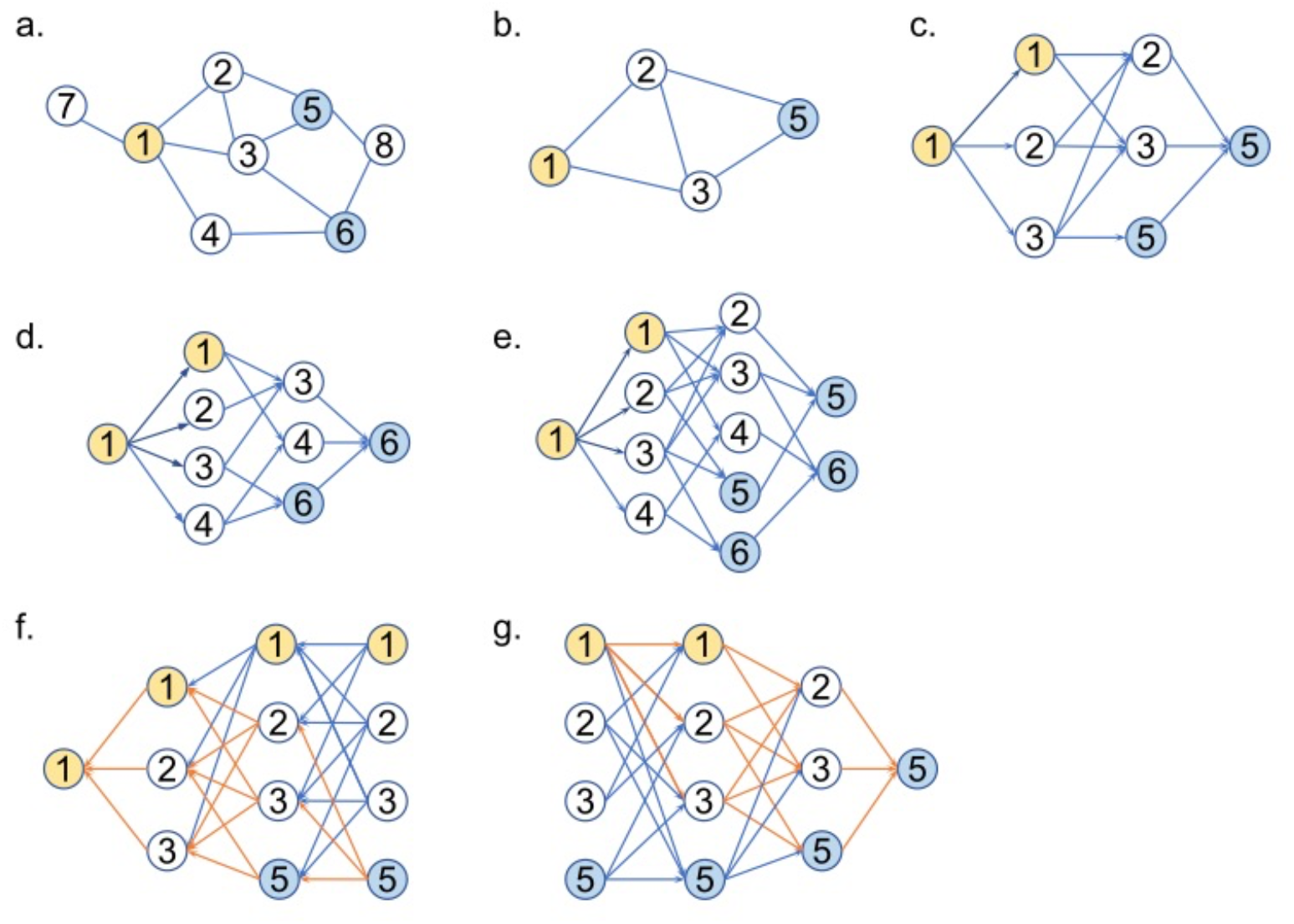
Illustration of the difference in message flow between RED-GNN/NBFNet and traditional GNN. (a) An example of KG; given query:(➀,r,?), answer:➄➅. (b) Subgraph for query (➀, r, ➄) with a path length of 3. (c)(d) The message passing flow for query (➀,r,➄), (➀,r,➅), respectively, in RED-GNN/NBFNet. (e) The message passing flow in (c) and (d) can be done in parallel. (f)(g) In the case of query (➀,r,➄), the message passing flow for head and tail in traditional GNN, respectively. The orange arrows indicate all the edges appearing in (c).

In the context of GraIL [23], the subgraph was trained as in a traditional GNN method, where each node aggregates its k-th order neighbor features, and the computation of *h*_*q*_ (*u, v*) goes for the mean node representation underlying the subgraph. It was suggested by the two following methods that the mean node representation may contain noise, as it is important to capture the significance of individual nodes only.

RED-GNN [25] addresses this issue from a message passing perspective, by utilizing the head node as the singular source node and the tail node as the singular target node. The message is transmitted from the sole source node to its 1-hop neighbor, where the 1-hop neighbor aggregates the message and subsequently transmits the new message to the 2-hop neighbors. This process continues, with each subsequent neighbor aggregating and transmitting the message to the next neighbor, until it reaches the target node after k steps.

For the example in Figure 2(b), the message passing flow is shown in Figure 2(c), and similar result can be obtained for the example of *h*_*q*_ (➀, ➅) as in Figure 2(d). Such process can be done in parallel (Figure 2(e)). Finally, the tail representation will be taken as the subgraph representation corresponding to *h*_*q*_ (*u, v*). The representation formula in RED-GNN can be articulated as follows:

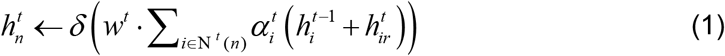

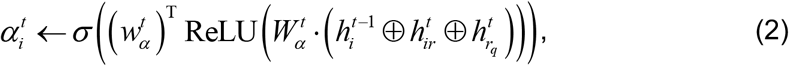

where 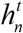 is the node representation in the t-th layer, 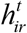 is the relation representation from node *i* to node *n* in the t-th layer, *w*^*t*^ ∈ ^*d*×*d*^ is a weighting matrix,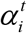is the attention score from node *i* to node 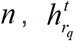 is the query representation in the t-th layer,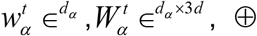 is the concatenation operator, N ^*t*^ (*n*)is the neighbor nodes passing message to node *n* in the t-th layer, *σ* is the Sigmoid function and *δ* is the nonlinear activation function.

NBFNet [24] aims to establish paired representations *h*_*q*_ (*u, v*) for all paths in the network. These paired representations *h*_*q*_ (*u, v*), which commence at a specific node *u* and conclude at another node *v*, are formulated as a generalized sum of path representations. Each path representation is characterized as a generalized product of the edge representations within the path, utilizing the multiplication operator:

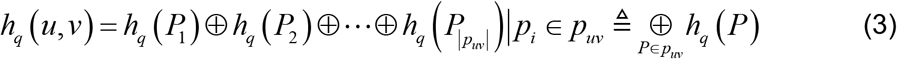

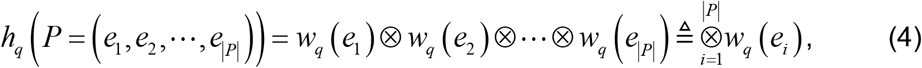

where *h*_*q*_ (*P*_*i*_) stand for the representation of path *i* under query *q*, ⊕ represents an arbitrary abstract function for aggregating paths without considering their order, *w*_*q*_ (*e*_*i*_) denotes for the representation of edge *i* under query *q*, ⊗ represents an abstract form of multiplication.

Unlike conventional multiplication, it does not necessarily adhere to the commutative property. For instance, such as matrix multiplication. To aggregate all the path messages, Depth-First Search can be used, but it becomes computationally challenging as the number of paths grows exponentially with the path length. NBFNet [24] proposed that if the abstract operations (⊕, ⊗) form a semiring, this problem can be transformed into an efficient computation using Breadth-First Search by General Bellman-Ford iteration:

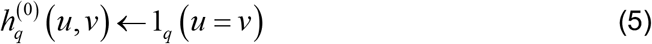

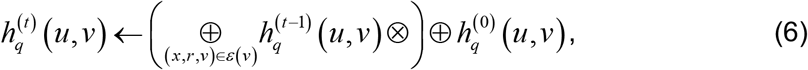

where 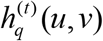 denote the representation in t-th step. 1_*q*_ (*u* = *v*) Denote the initial represents.

Inspire by formula (5) and formula (6), NBFNet relax the semiring assumption and incorporates the utilization of three neural networks to derive the following equations:

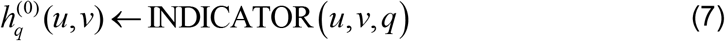

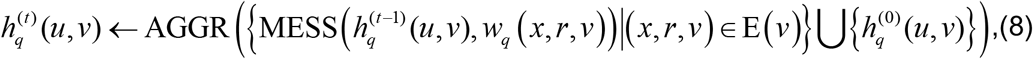

where INDICATOR, AGGR, MESS are all Neural Networks. Upon comparing formula (1) with formula (8), it is observed that both adhere to the message passing paradigm of GNN. Formula (8) exhibits a higher level of abstraction compared to formula (1)’s more specific nature. Consequently, they can be considered equivalent.

It is challenging to design a neural network that conforms to the semiring due to the presence of non-linear activation functions. As a result, the representation obtained from formula (8) does not actually rely on the path, but rather reverts to a representation based on subgraphs, which is equivalent to formula (1).

In traditional GNN, since each node aggregates its k-th order neighbor features, too many noisy nodes were introduced as shown in Figure 2(f, g). In contrast, The message passing flow of RED-GNN or NBFNet retains only the message from the head node ➀, making it a source-specific model, as illustrated in Figure 2(c). In other words, the representation of the tail relies exclusively on the message sending from the head node as the source, while in the traditional GNN all nodes become as the message passing flow source.

An effective method for achieving interpretability involves identifying the top-k significant edges. RED-GNN [25] employs attention to convey the significance of edges, whereas NBFNet [24] uses the partial derivatives of the scoring function to characterize the importance of each edge.

## RESULTS

### Construction of MegaKG

MegaKG integrated a total of 23 primary data sources, and finally consisted of 188, 844 nodes/entities and 9, 165, 855 edges/relations after processing. The entities were classified into 7 categories. In addition to the three fundamental categories including gene, disease and drug, there were also anatomy, phenotype, pathway and BP/MF/CC (ontology) which were affiliated to one or more of the fundamental categories. Relations involving one or two of the above mentioned entity categories, correspondingly, went into 20 major types. Please refer to Supplementary Table 1 and 2 for the summary of the entities and relations, respectively.

### Model performance evaluation

In our dataset, only gene-gene interaction relations are symmetric. Following the recommendation of NBFNet [24], we set the reverse relations of those symmetric relations to be the same as the original ones. We set the path length to be 4, the dimension of the relation embeddings to be 32, and performed negative sampling with 32 samples for each positive sample. Despite training for only one epoch, the model had converged, and its performance on the validation set was not improving.

We compared the performance of NBFNet [24] with other models on our dataset, and chose mean rank (MR) [10], mean reciprocal rank (MRR) [10], HITS@N [10] as the metrics. Notably as shown in Table 1, NBFNet outperformed other models in all metrics except MR.

### Case studies

To demonstrate the practical usage of MegaKG, especially for the early drug development in pharmaceutical industry, case studies were conducted. We focused on three typical scenarios - drug target prediction, indication extension and drug repurposing. In all cases, we prioritized the predictions based on the scores computed on the pair representation, and kept only those linkages whose corresponding relations were not present in the primary data sources underlying MegaKG. Uncollected evidences including, but not limited to, news report, cooperation announcements and scientific publications were leveraged to establish the credibility of these predictions. More importantly, for each predication, the explainable paths were provided, which may greatly help to judge the feasibility of the hypothesis or design downstream validation experiments, making deep learning techniques more than just black-box models for pharmaceutics.

*Case 1: Drug target prediction for Parkinson’s disease (PD)*. Supplemental Table 3 lists the top 50 candidate gene targets for treating PD whose disease-target relations were not collected in the primary data sources. We chose SORT1 as the demo case in Figure 3a, whose potential of such a relation was enhanced in the clinical trials of Humira [45]. Figure 3a also demonstrates the rationales behind to derive such a hypothesis. The largest contribution to link SORT1 to PD came from the protein-protein interaction with NGFR and BIN3. At the same time, essential tremor performed as a core node to form an alternative major explanatory route to PD. The relation of essential tremor and SORT1 has been recorded in the DrugBank [1]. Clues on this alternative root mainly included multiple genes involved with the pathogenesis of essential tremor.

**Figure 3.**
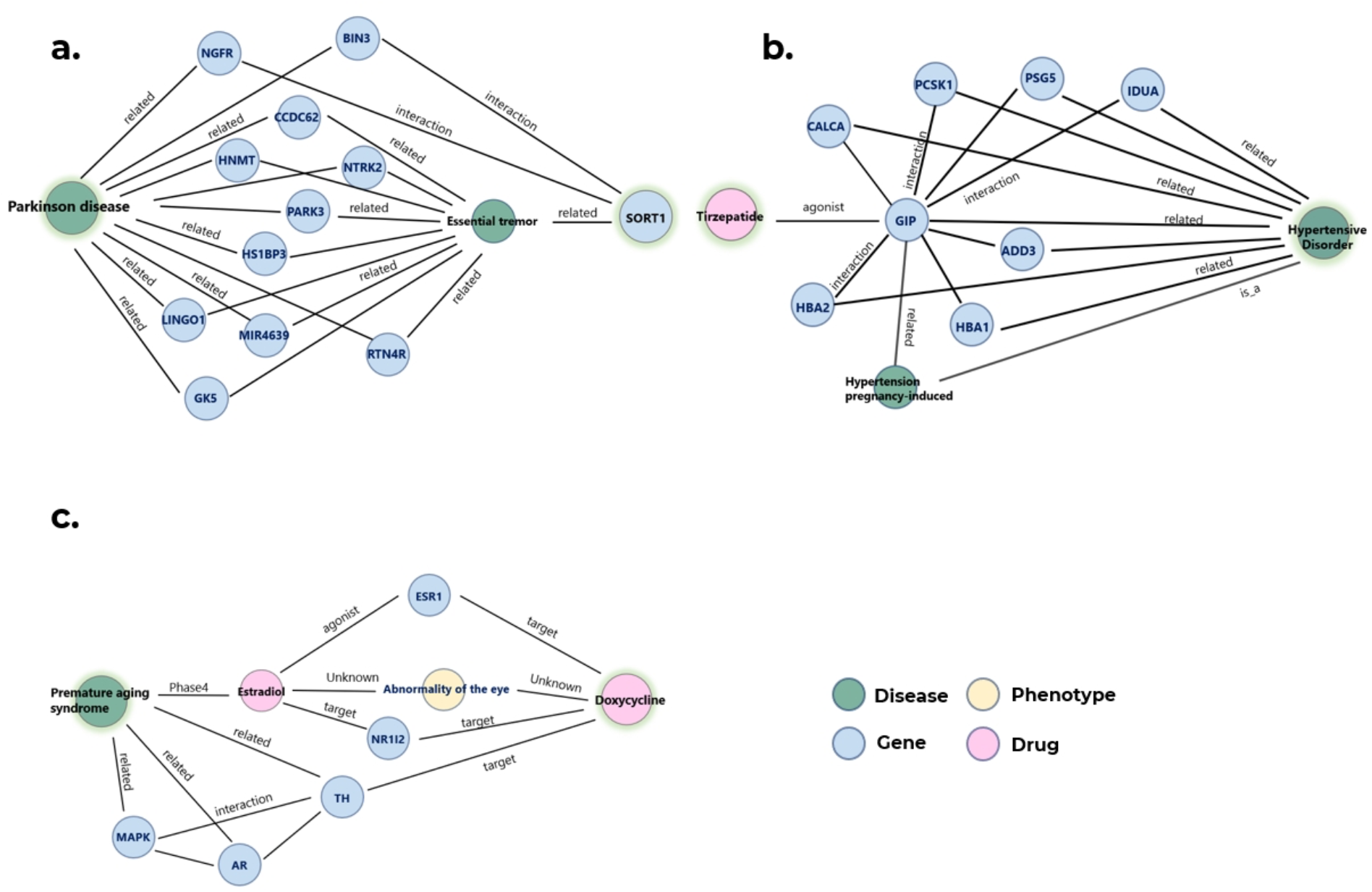
Explainable predictions for (a) Drug target prediction for Parkinson’s disease (PD); (b). Indication extension for Tirzepatide; (c) Drug-repurposing for premature aging syndrome.

*Case 2: Indication extension for Tirzepatide*. Tirzepatide, a dual glucose-dependent insulinotropic polypeptide (GIP) and glucagon-like peptide-1 (GLP-1) receptor agonist, has gained significant popularity in recent times and became a strong competitor of Semaglutide [1, 46]. Great marketing value may hold to extend the indication for such a drug. Therefore, we made predictions using MegaKG to relate Tirzepatide to other diseases not collected in the primary data sources, and a list of top 50 candidates were shown in Supplemental Table 4. A notable hypothesis was the Tirzepatide to treat hypertension, which was confirmed by the announcement of de Lamos et al [47]. As displayed in Figure 3b, most of the clues were through the GIP gene with its interacted genes between Tirzepatide and hypertension according to the initial records of DisGeNET [34]. The continuous connection from Tirzepatide to GIP and finally to hypertension constructed convincible interpretation to such a link prediction.

*Case3: Drug-repurposing for premature aging syndrome*. Premature aging syndrome belongs to a group of rare genetic disorders, for which available treatments are rather limited and specific drug development is slow. In such a situation, drug-repurposing is a practical approach. Doxycycline, a well-known nutritional product, appeared in the MegaKG predicted top 10 list (Supplemental Table 4), although its relation to any aging syndrome was not recorded in the databases [1, 46]. This drug repurposing potential was enhanced by the report of Bonuccelli et al [48]. One major explanation of the proposed connection was through the node of estradiol between Doxycycline and other node types as displayed in Figure 3c. And the contribution of estradiol to anti-aging was reported in Wise *et al*. [49].

Apparently, the predicted linkage from aging related drugs to Doxycycline was partly associated through nodes of phenotypes. Furthermore, another explainable path through genes like tyrosine hydroxylase (TH) has been highlighted, with facts collected in database [1, 45, 46].

### Comparison of prediction performance with known knowledge graphs

To further highlight the superior performance of MegaKG, two tasks were conducted in parallel following PharmKG (ref PharmKG), focusing on the two prevalent neurodegenerative disease -PD, and Alzheimer’s disease (AD). In the task of target gene prediction for Parkinson (Table 2A), 3 of the top 10 ranked coding genes for therapy inferred by MegaKG had been covered by drugs in clinical trials and 1 was in discovery state. In contrast, no genes predicted by PharmKG went into clinical trials, with only three genes in discovery state. In the other task of drug repurposing to AD, 6 of the top 10 drug predictions by MegaKG were in clinical trials, while only 1 was hit by PharmKG (Table 2B). It is evident to see that MegaKG exhibits superiority on the prediction efficacy than PharmKG, not to mention that it provides more reasonable interpretability as discussed in the Introduction.

**Table 2A:** Prediction comparison between PharmKG and MegaKG in the task of target gene prediction for PD.=.

| Alzheimer | PharmKG |  | MegaKG |  |
| --- | --- | --- | --- | --- |
|  | Drug | Clinical Trial Phase | Drug | Clinical Trial Phase |
|  | Etoposide | No | Pittsburgh Compound B | II |
|  | Midazolam | I | Acamprosate | II |
|  | Cyproterone acetate | No | Propofol | IV |
|  | Enalapril | No | Microcrystalline cellulose | II |
|  | Flupirtine | No | Oxygen | No |
|  | Imatinib | No | Methylinositol | No |
|  | Trifluoperazine | No | CX516 | II |
|  | Desipramine | No | Ezeprogind | No |
|  | Isoprenaline | No | MK 8719 | No |
|  | Imatinib | No | Piracetam | Un |

**Table 2B:** Prediction comparison between PharmKG and MegaKG in the task of drug repurposing to AD.

| Parkinson | PharmKG |  | MegaKG |  |
| --- | --- | --- | --- | --- |
|  | Gene | Clinical Trial Phase | Gene | Clinical Trial Phase |
|  | CYP3A4 | No | SLC17A7 | No |
|  | PSEN2 | No | TK2 | No |
|  | PSEN1 | No | KCNH4 | No |
|  | CYP2C8 | No | SORT1 | I |
|  | Sirtuin-2 | Discovery | SV2A | IV |
|  | TNFRSF1A | No | BIRC5 | No |
|  | ataxin-3 | Discovery | CALM3 | No |
|  | BIM | No | IL2 | I |
|  | ABCC2 | No | OPRK1 | Discovery |
|  | ataxin-3 | Discovery | SYNPO | No |

## DISCUSSION

In this study, we constructed a comprehensive biomedical KG, MegaKG, by integrating a total of 23 primary data sources, which finally consisted of 188, 844 nodes/entities and 9, 165, 855 edges/relations. Data within this field are diverse and disorganized, which lacks uniformity. We applied rigorous data processing criteria to guarantee high source quality to build MegaKG. In cases where the relations were ambiguous, we opted for a conservative approach to maintain accuracy. For example, drug-disease relation with both ‘contraindication’ and ‘indication’ from different database will be labeled as ‘unknown ‘. However, a lack of true negatives prevalently holds for all relation types inherently, to varying degrees. This poses a challenge to the sampling process to train a model to uncover hidden relations.

While several other biomedical KGs capable of predicting unknown relations have been developed, the majority of them could not provide explanations for their predictions properly. This brings a significant gap in the effective utilization of KGs in practical applications. PharmKG [7] moved a step forward by developing the HRGAT framework incorporating an attention mechanism to establish the interpretability for the results. However, such a framework can only provide static explanations in that once the network has finished training, the attention assigned to each node is fixed, irrespective of the specific queries. Adopting an approach based on pair representation, such as NBFNet [24], can successfully address this issue.

Most of the time, there exists a trade-off between scale and interpretability in models. NBFNet [24] exhibits a desirable characteristic: it achieves comparable performance even when trained on a mere 25% of the dataset, as compared to using the entire dataset. This finding suggests that the model’s demand for data is remarkably low and its performance remains consistent even with limited training data. This observation has significant implications for practical applications, especially in scenarios where data availability is limited or expensive. Researchers and practitioners can leverage this efficiency to build effective models even with minimal data, thus reducing computational costs and data collection efforts.

For the future extensive utilization of MegaKG, combining it with multi-omic analysis may emerge as a practical direction that can unlock its full value. For example, in the context of Mendelian disorders and rare-diseases, which affect millions of people each year, exome or genome sequencing have greatly facilitated prompt and accurate diagnoses. However, even after applying various filters, a typical sequencing result still contains hundreds of variants, necessitating significant time and clinical expertise to interpret them properly in relation to phenotypes and mutated genes. Tools incorporating MegaKG can be developed to prioritize genes by matching patient phenotypes. The combination of MegaKG and multi-omic data holds great promise in covering a wider range of available data modalities, and is anticipated to facilitate a wealth of discoveries.

## AUTHOR CONTRIBUTIONS

Q.W. conceived and initiated the study; J. D., J. L., and P. H. collected and preprocessed the primary data, and curated MegaKG; J.D. was in charge of link predication with emphasis on explainability, trained the models and performed evaluation results. Y. W. conducted case studies. All authors wrote and reviewed the manuscript.

### ACKNOWLEDGEMENTS

The authors would like to thank the other group members for helpful discussion and assistance.

